# EFFECT OF VISUAL-CUEING ON TWO-LEGGED HOPPING VARIABILITY IN CHILDREN WITH AUTISM SPECTRUM DISORDER: A PILOT STUDY

**DOI:** 10.1101/2020.02.05.936476

**Authors:** Daniel E. Lidstone, Janet S. Dufek

**Author notes:** Corresponding author: Daniel E. Lidstone, MS, Department of Kinesiology and Nutrition Sciences, University of Nevada, Las Vegas, 4505 S. Maryland Parkway Box 453034, Las Vegas, NV USA.

## Abstract

**Background:** Motor deficits in children with Autism Spectrum Disorder (ASD) are highly prevalent. High variability of motor output is commonly reported in children with ASD. Visual cueing using an exergame may be an effective intervention to reduce motor variability in children with ASD.

**Aim:** To examine the effect of visual cueing on two-legged hopping variability in children with ASD.

**Methods:** Four children with ASD and six age-matched TD controls performed three 20-s hopping trials with no visual cueing (no cue = NC) and with a 2 Hz visual cue (visual cue = VC). Three-dimensional kinematic data of the sacrum marker and ground reaction force were collected during each hopping trial. Variability was determined using the intra-trial coefficient of variation (CoV) of hopping frequency, hop height, and negative sacral displacement

**Results:** A marginally significant interaction between GROUP (ASD/TD) and CUE type (NC/VC) was observed for hopping frequency variability (*p* = 0.06) indicating greater impairment in the ASD group vs. TD group with visual vs. no-cueing. The main effect of group showed a statistically significant difference in hopping frequency (*p* = 0.037), hopping frequency variability (*p* = 0.008), and negative sacrum displacement variability (*p* = 0.04).

**Conclusion:** This pilot study confirmed high motor variability in the amplitude and frequency of repetitive movements in children with ASD. However, visual cueing was ineffective at reducing the variability of motor output in children with autism.

## 1. INTRODUCTION

Autism Spectrum Disorder (ASD) is a neurodevelopmental disorder characterized by social and communication impairments, restricted interests, and repetitive behaviors (APA, 2000). In addition to the core features of ASD, motor deficits are highly prevalent in children with ASD (Berkeley et al. 2001; Green et al. 2009; Fournier et al. 2010; Barrow et al. 2011; Liu 2014) with approximately 8 in 10 children with autism showing definite motor impairments (Green et al. 2009). Motor control impairments are one of the earliest signs of autism emerging as early as 7 months (Leonard et al. 2014), whereas core social communication deficits emerge later when infants are ∼2 years old.

Although the motor impairments observed in children with ASD are not solely due to intellectual ability (Rinehart et al. 2006; Green et al. 2009; Travers et al. 2013), 70% of persons with ASD have an intellectual disability (ID) (La Malfa et al. 2004) and these children show more severe motor impairments compared to children with ASD and co-occurring ID (Green et al. 2009). Therefore, there is a need to develop targeted movement interventions to improve motor skills in children with ASD that involve minimal instructions. Exergames that provide visual cueing are simple interventions, requiring minimal verbal instruction, that may help improve gross-motor control in children with ASD and children with ASD and co-occurring ID. Although visual feedback has been proposed as a strategy to improve motor control in children with ASD (Somogyi et al. 2016), visuomotor integration deficits are a relatively consistent finding in this population (Masterton and Biederman 1983; Haswell et al. 2009; Dowd et al. 2012; Greffou et al. 2012; Papadopoulos et al. 2012; Ament et al. 2015; Nebel et al. 2016; Sharer et al. 2016). Furthermore, there is evidence that children with ASD show impaired integration of visual cueing and feedback to modulate ongoing fine-motor (Mosconi et al. 2015; Wang et al. 2015, 2017) and gross-motor commands (Nayate et al. 2012), with increased visuomotor integration demands associated with increased variability of motor output compared to TD children. Individuals with ASD also show deficits during whole-body gross motor tasks involving multi-modal integration of auditory and proprioception sources (Moran et al. 2013). However, the effects of visual cueing via an exergame on gross-motor control has not been examined in children with ASD.

In the current study, we examined the effects of whole-body multi-modal integration of sensory input from visual and proprioception sources. In similarity to Moran et al. (2013), we chose to cue the timing of a repetitive gross-motor task using a fixed cadence stimulus, however, instead of using a metronome auditory cue, a 2 Hz visual rhythmic cue was used. The purpose of this exploratory study was to examine the effect of an exergame with visual cueing on movement variability in children with autism. Because previous studies have documented increased gross motor variability in children with ASD, we first hypothesized that (1) children with ASD will be more variable in frequency and amplitude of hopping compared to TD children. Second, because of well documented impairments in visuomotor integration in children with ASD we hypothesized that (2) children with ASD, but not TD children, will have significantly greater hopping variability in frequency and amplitude of hopping during visual cueing (VC) compared to no-cue (NC) conditions.

## 2. MATERIAL AND METHODS

### 2.1 Participants

Seven children (10-15 years old) with parent-confirmed medical diagnoses of ASD and seven age-matched TD children without medical diagnoses of any neurological disorder were recruited for participation in the study (Table 1). Participants were well matched for age, body mass, and height (Table 1). All parents confirmed diagnoses and that their child had: (1) normal (20/20) or corrected-to-normal vision (20/20), (2) no history of lower limb surgery, and (3) no current orthopedic injury. The presence of comorbid psychiatric disorders was not formally assessed in either group. Three children in the ASD group and one child in the TD group did not complete the study due to behavioral issues. Two of the children in the ASD group that did not complete the study were non-verbal. Institutionally approved parental consent and child assent were obtained prior to participation.

**Table 1:** Participant demographics. Significant differences between groups for age, body mass, and height performed using an independent samples t-test. A priori alpha was set at 0.05.

|  | **ASD (n=4)** | **TD (n=6)** | ***p* value** |
| --- | --- | --- | --- |
| **Sex (M/F)** | 3/1 | 5/1 | - |
| **Age (years)** | 12.2±2.5 | 12.2±1.8 | 0.989 |
| **Body Mass (kg)** | 45.7±13.6 | 47.5±10.9 | 0.827 |
| **Height (m)** | 1.56±0.2 | 1.51±0.1 | 0.604 |

### 2.2 Instrumentation

Three dimensional (3D) kinematic data from a retro-reflective marker on the sacrum of each child was obtained using a 10-camera motion capture system (100 Hz; Vicon Motion Systems, Ltd.). Simultaneous kinetic data were obtained using two Kistler force platforms (1000 Hz; Kistler, Amherst, NY) that were mounted flush with the floor and positioned side-by-side.

### 2.3 Visual Stimuli

The visual stimuli (2 Hz bouncing ball) used in the VC condition was presented using a custom-built exergame (Idoneus Digital®, Vancouver, Canada) developed in Unity™. A frequency of 2 Hz was selected since it is close to the preferred frequency of human hopping where the center of mass behaves like a spring-mass system (Blickhan 1989). The bouncing ball behaved as a perfectly elastic object under the influence of gravity, returning to the same height with each bounce. The game did not produce any sound, thereby only providing visual stimuli.

### 2.4 Procedure

During the experiment the children wore tight-fitting spandex shorts and their own athletic shoes. The two-legged hopping task was demonstrated by the experimenter standing in front of the child, however, the children were not given any instructions on how high to hop (Moran et al. 2013). To reduce the influence of aberrant arm movements during the hopping task, the children performed the hopping task with their arms folded across their chest (Figure 1). A retro-reflective marker was affixed to the sacrum using double-sided and Cover-Roll Stretch® tape. Participants were then positioned so they were standing on both force plates. The force plates were 1.5 meters from a television (55 in LED display, 3840×2160 resolution, 120 Hz refresh rate, Polaroid, MN, USA) monitor that was mounted on a height-adjustable TV stand. The TV height was adjusted so the center of the screen was set approximately at eye-level (Figure 1).

**Figure 1:**
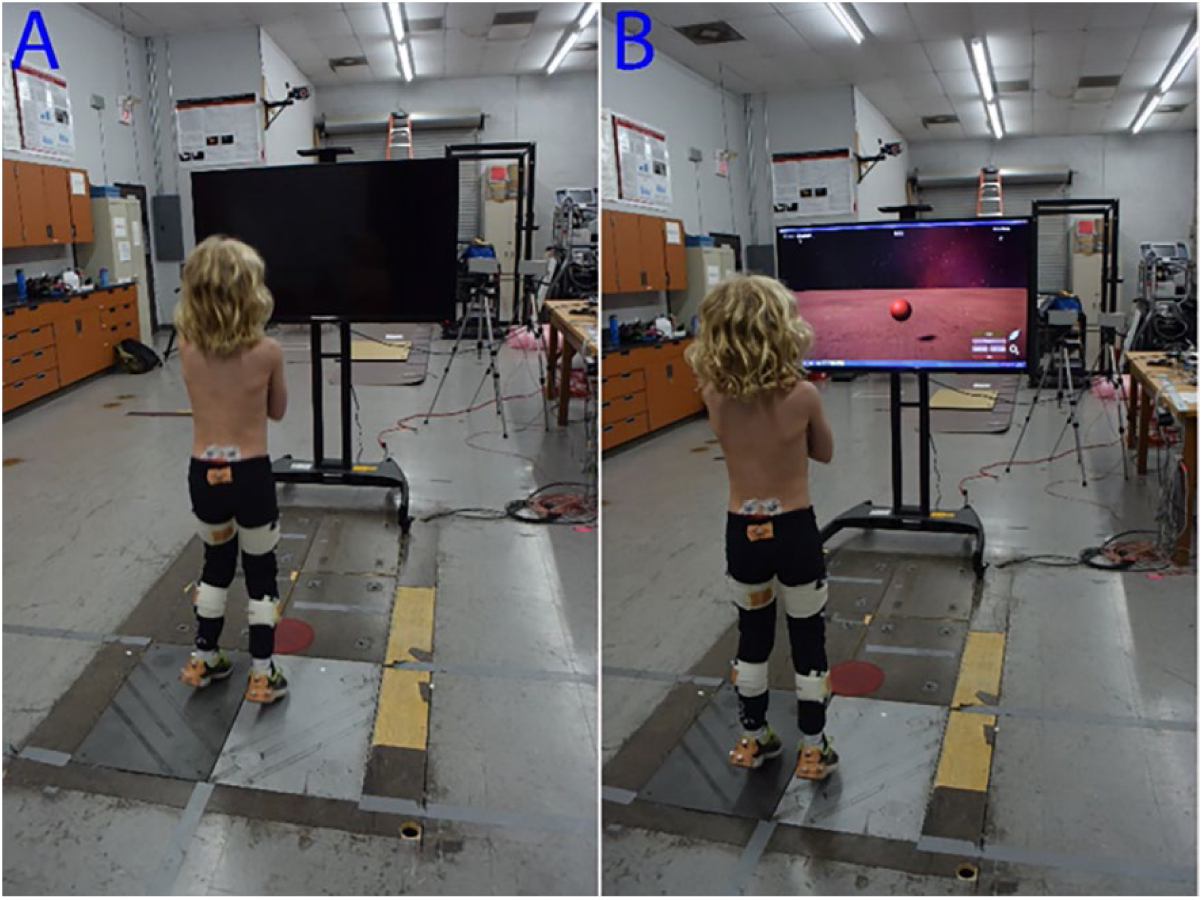
Experimental set-up. A = no cue condition (NC); B = 2 Hz visual cue condition (VC).

Each child performed three 20 second hopping trials in two conditions: (1) NC and (2) VC. At the beginning of each condition, the participants completed one full 20-second practice trial to ensure they understood the task. The NC condition was always performed first to reduce the effects of visual cueing on the self-selected hopping pattern.

In the NC condition the children hopped at a self-selected height and frequency for 20 seconds. In the VC condition, the children were instructed to hop in unison with the 2 Hz bouncing ball. Each trial lasted 20 seconds and two-minutes of rest were provided between trials. Five minutes of rest were provided between conditions.

### 2.5 Post-Processing Procedures

All post-processing procedures were performed using custom scripts in Matlab software (Matlab 2017, Natick, USA). First, kinetic data were down sampled to 100 Hz to match the sampling frequency of kinematic data. Sacrum marker trajectories and kinetic data were filtered using a 4^th^ order low-pass Butterworth filter with cut-offs of 6 Hz and 25 Hz, respectively. A 25 Newton (N) cut-off was then used to identify ground contact (GC) and toe-off (TO) for each hop. The first two hops of each trial were not included in the analysis with hops 3 to 27 selected for processing (Figure 2). This number of consecutive hops were selected since each participant performed at least 25 consecutive hops, without stopping, in each trial.

**Figure 2:**
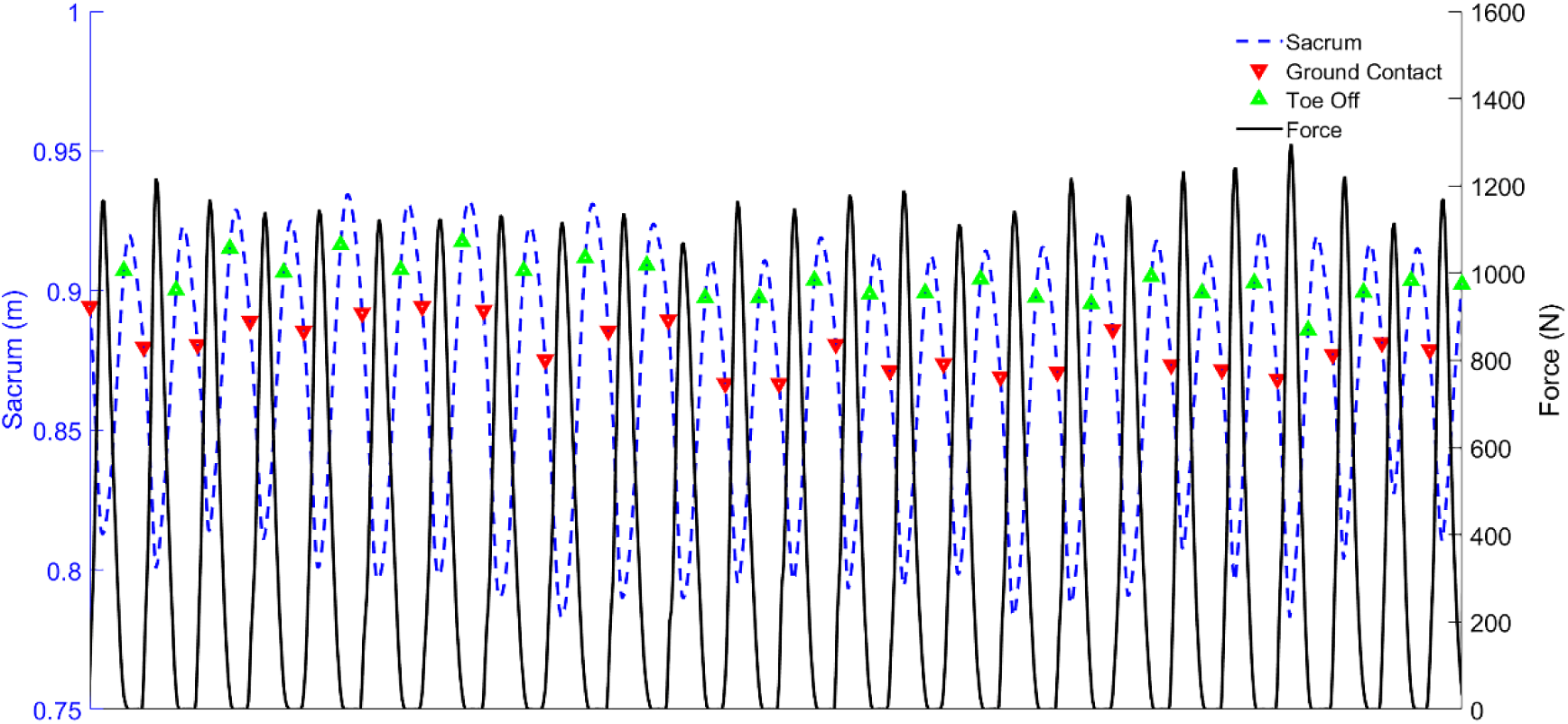
Sacrum marker, ground-reaction force (GRF) data, and event detection (ground contact and toe-off) for 25 consecutive hops during a single trial.

### 2.6 Hop Frequency and Amplitude Variability

Hop frequency (Hz) was calculated as the inverse of time in seconds (T) from GC to the subsequent GC (*f*=1/T). The mean hop frequency and coefficient of variation (CoV) [CoV = (SD/Mean)100] for each trial was calculated using the 25 consecutive hops within each trial. Two measures of sacrum amplitude variability were examined for all hops within each trial: (1) absolute change in sacral marker amplitude from GC to minimum sacrum position (negative sacrum displacement; NSD) and (2) absolute change in sacral marker amplitude from TO to maximal sacrum height (hop height; HH) (Figure 2). The CoV was calculated, using 25 consecutive hops within each trial, for NSD and HH.

### 2.7 Statistical Procedures

All statistical analyses were performed using SPSS 24 (IBM Corp., Armonk, NY). A two-way mixed ANOVA was performed with two between subject’s factors (ASD and TD) and two within subject’s factors (NC and VC). All data were assessed for normal distribution using Shapiro-Wilk’s test of normality and assumptions of homogeneity of variances met as determined by Levene’s test. An *a priori* level of significance was set at 0.05. Cohen’s *d* effect sizes (ES) were also calculated for each dependent variable within each condition [Cohen’s *d* = (*M*_*ASD*_− *M*_*TD*_)/*SD*_*pooled*_]. Large ES were defined as ES>0.80 (Cohen 1988).

## 3. RESULTS

There were no statistically significant interactions between group and cue type for hopping frequency (*F*_(1,8)_ = 1.535, *p* = 0.251, n^2^ = 0.161), hopping frequency variability (*F*_(1,8)_ = 4.607, *p* = 0.064, n^2^ = 0.365), negative sacrum displacement variability (*F*_(1,8)_ = 0.144, *p* = 0.714, n^2^ = 0.018), and hop height variability (*F*_(1,8)_ = 0.177, *p* = 0.685, n^2^ = 0.022) (Table 2). However, the group x cue interaction for hop frequency variability was marginally significant at *p* = 0.06, with the ASD group showing greater hopping variability compared to the TD group in the VC compared to the NC condition (Figure 3). The main effect of group showed a statistically significant difference in hopping frequency (*F*_(1,8)_ = 6.248, *p* = 0.037, n^2^ = 0.439, ASD>TD), hopping frequency variability (*F*_(1,8)_ =, *p* = 0.008, n^2^ = 0.604, ASD>TD), and negative sacrum displacement variability (*F*_(1,8)_ = 5.977, *p* = 0.04, n^2^ = 0.428, ASD>TD). Effect sizes were large (ES>0.80) for all dependent variables except for hop height variability in the NC and VC conditions that showed moderate ES (moderate ES = 0.51-0.80). All ES were positive showing that children in the ASD hopped at a faster cadence and were more variable in hopping frequency and amplitude in each condition (Table 2).

**Table 2:** Summary of results of two-way mixed ANOVAs and effect sizes. Significant p values and large effect size’s (ES>0.80) in bold.

|  |  | ASD | TD | ES | Two-Way Mixed ANOVA |  |  |
| --- | --- | --- | --- | --- | --- | --- | --- |
|  |  |  |  |  | Cue*<br>Group | Cue | Group |
| Hopping<br>Frequency<br>(Hz) | NC | 2.49±0.42 | 2.02±0.16 | **1.47** | 0.251 | 0.933 | **0.037** |
|  | VC | 2.42±0.35 | 2.10±0.10 | **1.24** |  |  |  |
| Hopping<br>Frequency<br>(CoV - %) | NC | 5.63±1.84 | 3.99±0.86 | **1.14** | 0.064 | 0.076 | **0.008** |
|  | VC | 7.31±1.90 | 3.95±0.43 | **2.43** |  |  |  |
| Negative Sacrum<br>Displacement<br>(CoV - %) | NC | 15.41±5.05 | 10.08±1.93 | **1.39** | 0.714 | 0.179 | **0.04** |
|  | VC | 16.20±5.01 | 11.42±1.64 | **1.28** |  |  |  |
| Hop Height<br>(CoV - %) | NC | 25.04±8.56 | 19.67±5.35 | 0.75 | 0.685 | 0.184 | 0.187 |
|  | VC | 31.56±9.91 | 23.26±11.55 | 0.77 |  |  |  |

**Figure 3:**
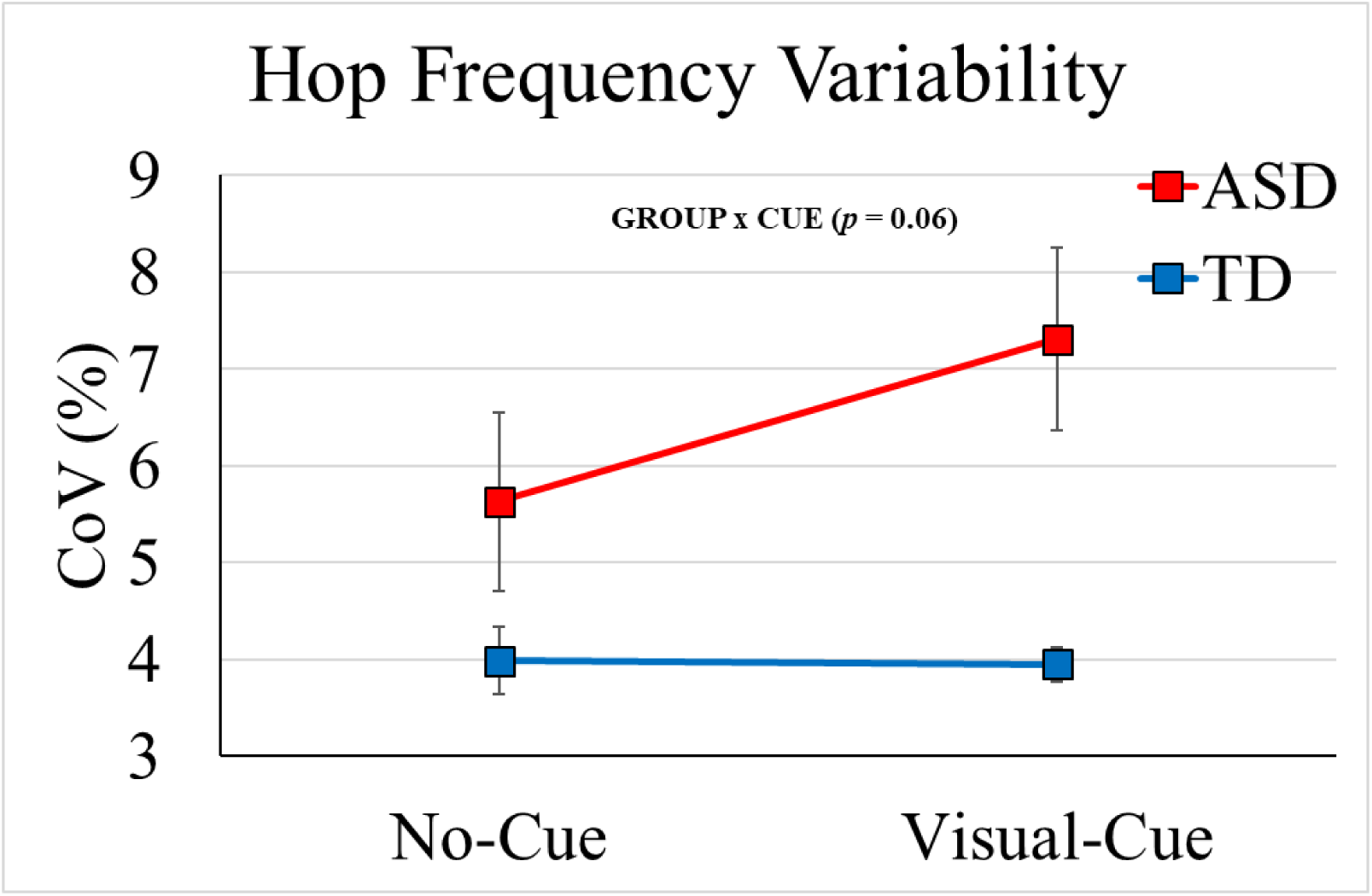
Hop frequency variability (CoV %) means and standard errors for ASD and TD groups during no-cue (NC) and visual-cue (VC) conditions.

## 4. DISCUSSION

The purpose of this exploratory study was to determine the effect of visual cueing on gross motor movement variability in children with ASD. The main findings of this exploratory study were: (1) children with ASD were more variable in both hopping frequency and amplitude than TD children, in agreement with our hypothesis, and (2) visual cueing had no significant effect on hopping variability in frequency or amplitude in either group, in rejection of our hypothesis. However, a marginally significant group x cue interaction was observed for hopping frequency variability (*p* = 0.06) that suggested increased variability in the ASD vs. TD group with visual cueing compared to without visual cueing. Therefore, dynamic visual cueing as used in this study may not be effective at decreasing gross motor variability in children with autism. No other study to date has examined the effect of an exergame, using dynamic visual cueing, on the variability of a gross-motor task in children with ASD.

The high variability of motor output in the ASD during the visual cueing condition indicates that the children in the ASD group had difficulties integrating visual input with motor output (Parma and de Marchena 2016). Furthermore, children with ASD were unable to adequately match the 2 Hz visual cue. In the VC conditions, the ASD group deviated 21% from the 2 Hz visual cue, whereas the TD group deviated only 5%. However, the self-selected cadence of the TD group was closer to the 2 Hz cue (1% faster) compared to the ASD group (24.5% faster) and therefore motor commands would have been similar between conditions for the TD group. Future studies should use a wider range of frequencies to examine whether our observation that children with ASD are unable to match a dynamic visual cue are frequency dependent or specific to the visual and/or dynamic nature of the cue.

To our knowledge, only one other study has examined the effects of visual cueing on the variability of a gross motor task (Nayate et al. 2012). Nayate et al. (2012), examined the effect of visual spatial cueing on gait parameters in children with ASD and TD controls and found increases in stride length variability in the cued vs. un-cued conditions only in the ASD group. Our findings somewhat agree with Nayate et al. (2012) since we observed no improvements in motor control in children with ASD using visual cueing, however, we did not observe significant increase in movement variability with visual cueing. Our findings may be explained by atypical multi-modal integration of visual and proprioception sensory input as previously demonstrated in persons with ASD (Cascio et al. 2012). This may result from an overreliance on proprioception over visual sensory input (Izawa et al. 2012; Marko et al. 2015). Furthermore, deficits in tracking visual stimuli may also contribute to the ineffectiveness of visual cueing in altering the timing and variability of gross-motor output in children with ASD (Takarae et al. 2004).

In addition to visuomotor integration impairments in children with autism (Masterton and Biederman 1983; Inui et al. 1995; Gepner and Mestre 2002; Dowd et al. 2012; Greffou et al. 2012; Marko et al. 2015; Mosconi et al. 2015), oculomotor deficits are also widely reported (Takarae et al. 2004, 2007, 2014; Wegiel et al. 2013; Freedman and Foxe 2017). Deficits in both dynamic visual tracking and integration of visual input to update motor commands in children with ASD implicate the oculomotor and sensorimotor cerebellum. Cerebellum lobule VI is part of the oculomotor vermis and is consistently implicated in visuomotor learning tasks (Vaillancourt et al. 2006; Donchin et al. 2012; Neely et al. 2013; Moulton et al. 2017) and both structural and functional deficits of cerebellum lobule VI are consistently found in individuals with ASD (Courchesne et al. 1988, 1994; Skefos et al. 2014). Furthermore, cerebellum lobule VI is implicated in anomalous autism-associated proprioception bias (Marko et al. 2015) and the severity of core ASD symptoms (Kaufmann et al. 2003; Rojas et al. 2006; Takarae et al. 2007). Therefore, cerebellar structural and functional deficits in children with autism may limit the effectiveness of visual cueing stimuli to decrease gross motor variability.

### 4.1 Conclusions

In conclusion, this pilot study confirmed previous findings of high variability of motor output and supports visuomotor integration deficits in children with ASD. These findings have implications for the design of technological interventions to improve motor control in children with ASD. To the best of our knowledge, this is the first study to use biomechanics methods to examine the effect of an exergame on gross-motor variability in children with ASD. Future studies examining the effect of exergames on motor control in children with ASD should employ a larger sample size, control for IQ, control for ASD symptom severity, and utilize eye-tracking technology.

## Acknowledgements

None

## REFERENCES

Ament K, Mejia A, Buhlman R, Erklin S, Caffo B, Mostofsky S, et al. Evidence for Specificity of Motor Impairments in Catching and Balance in Children with Autism. J Autism Dev Disord. 2015;45(3):742–51.

American Psychiatric Association. DSM-IV-TR Diagnostic and statistical manual of mental disorders. 4th edn. American Psychiatric Association, Washington, DC; 2000.

Barrow WJ, Jaworski M, Accardo PJ. Persistent Toe Walking in Autism. J Child Neurol [Internet]. 2011;26(5):619–21. Available from: http://jcn.sagepub.com/cgi/doi/10.1177/0883073810385344

Berkeley SL, Zittel LL, Pitney L V., Nichols SE. Locomotor and object control skills of children diagnosed with autism. Adapt Phys Act Q. 2001;18(4):405–16.

Blickhan R. The spring-mass model for running and hopping. Vol. 22, Journal of Biomechanics. 1989. p. 1217–27.

Cascio CJ, Burnette CP, Cosby AA. The rubber hand illusion in children with autism spectrum disorders: delayed influence of combined tactile and visual input on proprioception. Autism. 2012;16(4):406–19.

Cohen J. Statistical power analysis for the behavioral sciences. 2nd. 1988.

Courchesne E, Saitoh O, Yeung-Courchesne R, Press GA, Lincoln AJ, Haas RH, et al. Abnormality of cerebellar vermian lobules VI and VII in patients with infantile autism: Identification of hypoplastic and hyperplastic subgroups with MR imaging. Am J Roentgenol. 1994;162(1):123–30.

Courchesne E, Yeung-Courchesne R, Hesselink JR, Jernigan TL. Hypoplasia of Cerebellar Vermal Lobules VI and VII in Autism. N Engl J Med [Internet]. 1988;318(21):1349–54. Available from: http://www.nejm.org/doi/abs/10.1056/NEJM198805263182102

Donchin O, Rabe K, Diedrichsen J, Lally N, Schoch B, Gizewski ER, et al. Cerebellar regions involved in adaptation to force field and visuomotor perturbation. J Neurophysiol [Internet]. 2012;107(1):134–47. Available from: http://jn.physiology.org/cgi/doi/10.1152/jn.00007.2011

Dowd AM, McGinley JL, Taffe JR, Rinehart NJ. Do planning and visual integration difficulties underpin motor dysfunction in autism? A kinematic study of young children with autism. J Autism Dev Disord. 2012;42(8):1539–48.

Fournier KA, Hass CJ, Naik SK, Lodha N, Cauraugh JH. Motor coordination in autism spectrum disorders: A synthesis and meta-analysis. J Autism Dev Disord. 2010;40(10):1227–40.

Freedman EG, Foxe JJ. Eye movements, sensorimotor adaptation and cerebellar-dependent learning in autism: Toward potential biomarkers and subphenotypes. Eur J Neurosci. 2017;1–7.

Gepner B, Mestre DR. Brief Report: Postural Reactivity to Fast Visual Motion Differentiates Autistic from Children with Asperger Syndrome. J Autism Dev Disord. 2002;32(3):231–8.

Green D, Charman T, Pickles A, Chandler S, Loucas T, Simonoff E, et al. Impairment in movement skills of children with autistic spectrum disorders. Dev Med Child Neurol. 2009;51(4):311–6.

Greffou S, Bertone A, Hahler EM, Hanssens JM, Mottron L, Faubert J. Postural hypo-reactivity in autism is contingent on development and visual environment: A fully immersive virtual reality study. J Autism Dev Disord. 2012;42(6):961–70.

Haswell CC, Izawa J, R Dowell L, H Mostofsky S, Shadmehr R. Representation of internal models of action in the autistic brain. Nat Neurosci. 2009;12(8):970–2.

Inui N, Yamanishi M, Tada S. Simple Reaction Times and Timing of Serial Reactions of Adolescents with Mental Retardation, Autism, and down Syndrome. Percept Mot Skills [Internet]. 1995;81(3):739–45. Available from: http://journals.sagepub.com/doi/10.2466/pms.1995.81.3.739

Izawa J, Pekny SE, Marko MK, Haswell CC, Shadmehr R, Mostofsky SH. Motor learning relies on integrated sensory inputs in ADHD, but over-selectively on proprioception in autism spectrum conditions. Autism Res. 2012;5(2):124–36.

Kaufmann WE, Cooper KL, Mostofsky SH, Capone GT, Kates WR, Newschaffer CJ, et al. Specificity of cerebellar vermian abnormalities in autism: A quantitative magnetic resonance imaging study. J Child Neurol. 2003;18(7):463–70.

Leonard HC, Elsabbagh M, Hill EL. Early and persistent motor difficulties in infants at-risk of developing autism spectrum disorder: A prospective study [Internet]. Vol. 11, European Journal of Developmental Psychology. Taylor & Francis; 2014. p. 18–35. Available from: http://dx.doi.org/10.1080/17405629.2013.801626

Liu T. Gross Motor Performance by Children with Autism Spectrum Disorder and Typically Developing Children on TGMD-2. J Child Adolesc Behav [Internet]. 2014;02(01). Available from: http://www.esciencecentral.org/journals/gross-motor-performance-by-children-with-autism-spectrum-disorder-and-typically-2375-4494.1000123.php?aid=22538

La Malfa G, Lassi S, Bertelli M, Salvini R, Placidi GF. Autism and intellectual disability: A study of prevalence on a sample of the Italian population. J Intellect Disabil Res. 2004;48(3):262–7.

Marko MK, Crocetti D, Hulst T, Donchin O, Shadmehr R, Mostofsky SH. Behavioural and neural basis of anomalous motor learning in children with autism. Brain. 2015;138(3):784–97.

Masterton BA, Biederman GB. Proprioceptive versus visual control in autistic children. J Autism Dev Disord. 1983;13(2):141–52.

Moran MF, Foley JT, Parker ME, Weiss MJ. Two-legged hopping in autism spectrum disorders. Front Integr Neurosci [Internet]. 2013;7(March):14. Available from: http://journal.frontiersin.org/article/10.3389/fnint.2013.00014/abstract

Mosconi MW, Mohanty S, Greene RK, Cook EH, Vaillancourt DE, Sweeney JA. Feedforward and Feedback Motor Control Abnormalities Implicate Cerebellar Dysfunctions in Autism Spectrum Disorder. J Neurosci [Internet]. 2015;35(5):2015–25. Available from: http://www.jneurosci.org/cgi/doi/10.1523/JNEUROSCI.2731-14.2015

Moulton E, Galléa C, Kemlin C, Valabregue R, Maier MA, Lindberg P, et al. Cerebello-Cortical Differences in Effective Connectivity of the Dominant and Non-dominant Hand during a Visuomotor Paradigm of Grip Force Control. Front Hum Neurosci [Internet]. 2017;11(October):1–12. Available from: http://journal.frontiersin.org/article/10.3389/fnhum.2017.00511/full

Nayate A, Tonge BJ, Bradshaw JL, McGinley JL, Iansek R, Rinehart NJ. Differentiation of high-functioning autism and Asperger’s disorder based on neuromotor behaviour. J Autism Dev Disord. 2012;42(5):707–17.

Nebel MB, Eloyan A, Nettles CA, Sweeney KL, Ament K, Ward RE, et al. Intrinsic visual-motor synchrony correlates with social deficits in autism. Biol Psychiatry. 2016;79(8):633–41.

Neely KA, Coombes SA, Planetta PJ, Vaillancourt DE. Segregated and overlapping neural circuits exist for the production of static and dynamic precision grip force. Hum Brain Mapp. 2013;34(3):698–712.

Papadopoulos N, McGinley J, Tonge BJ, Bradshaw JL, Saunders K, Rinehart NJ. An investigation of upper limb motor function in high functioning autism and Asperger’s disorder using a repetitive Fitts’ aiming task. Res Autism Spectr Disord [Internet]. 2012;6(1):286–92. Available from: http://dx.doi.org/10.1016/j.rasd.2011.05.010

Parma V, de Marchena AB. Motor signatures in autism spectrum disorder: the importance of variability. J Neurophysiol [Internet]. 2016;115(3):1081–4. Available from: http://jn.physiology.org/lookup/doi/10.1152/jn.00647.2015

Rinehart NJ, Tonge BJ, Iansek R, McGinley J, Brereton A V, Enticott PG, et al. Gait function in newly diagnosed children with autism: cerebellar and basal ganglia related motor disorder. Dev Med Child Neurol [Internet]. 2006;48(10):819. Available from: http://doi.wiley.com/10.1017/S0012162206001769

Rojas DC, Peterson E, Winterrowd E, Reite ML, Rogers SJ, Tregellas JR. Regional gray matter volumetric changes in autism associated with social and repetitive behavior symptoms. BMC Psychiatry. 2006;6:1–13.

Sharer EA, Mostofsky SH, Pascual-Leone A, Oberman LM. Isolating Visual and Proprioceptive Components of Motor Sequence Learning in ASD. Autism Res. 2016;9(5):563–9.

Skefos J, Cummings C, Enzer K, Holiday J, Weed K, Levy E, et al. Regional alterations in Purkinje cell density in patients with autism. PLoS One. 2014;9(2):1–12.

Somogyi E, Kapitány E, Kenyeres K, Donauer N, Fagard J, Kónya A. Visual feedback increases postural stability in children with autism spectrum disorder. Res Autism Spectr Disord. 2016;29–30:48–56.

Takarae Y, Luna B, Minshew NJ, Sweeney JA. Visual motion processing and visual sensorimotor control in autism. J Int Neuropsychol Soc [Internet]. 2014;20(1):113–22. Available from: http://www.journals.cambridge.org/abstract_S1355617713001203

Takarae Y, Minshew NJ, Luna B, Krisky CM, Sweeney JA. Pursuit eye movement deficits in autism. Brain. 2004;127(12):2584–94.

Takarae Y, Minshew NJ, Luna B, Sweeney JA. Atypical involvement of frontostriatal systems during sensorimotor control in autism. Psychiatry Res - Neuroimaging. 2007;156(2):117–27.

Travers BG, Powell PS, Klinger LG, Klinger MR. Motor difficulties in autism spectrum disorder: Linking symptom severity and postural stability. J Autism Dev Disord. 2013;43(7):1568–83.

Vaillancourt DE, Mayka MA, Corcos DM. Intermittent visuomotor processing in the human cerebellum, parietal cortex, and premotor cortex. J Neurophysiol. 2006;95(2):922–31.

Wang Z, Kwon M, Mohanty S, Schmitt LM, White SP, Christou EA, et al. Increased force variability is associated with altered modulation of the motorneuron pool activity in autism spectrum disorder (ASD). Int J Mol Sci. 2017;18(4).

Wang Z, Magnon GC, White SP, Greene RK, Vaillancourt DE, Mosconi MW. Individuals with autism spectrum disorder show abnormalities during initial and subsequent phases of precision gripping. J Neurophysiol [Internet]. 2015;113(7):1989–2001. Available from: http://jn.physiology.org/lookup/doi/10.1152/jn.00661.2014

Wegiel J, Kuchna I, Nowicki K, Imaki H, Wegiel J, Ma YS, et al. Contribution of olivofloccular circuitry developmental defects to atypical gaze in autism. Brain Res [Internet]. 2013;1512(2013):106–22. Available from: http://dx.doi.org/10.1016/j.brainres.2013.03.037

